# Adaptive feedforward speed control in *Drosophila*

**DOI:** 10.1101/2025.03.03.641162

**Authors:** Benjamin P. Campbell, Jack A. Supple, Samuel T. Fabian, Huai-Ti Lin, Holger G. Krapp

## Abstract

Insects demonstrate remarkable agility in flight despite constant changes in flight dynamics throughout their lives. However, it is unclear whether such resilience is conferred via purely feedback control or whether adaptive feedforward control mechanisms are present. This study examines whether adaptive feedforward control mechanisms are present in *Drosophila melanogaster* flight, by comparing the free-flight trajectories with and without wing damage and antennae ablation. Flies with partial wing excisions exhibited increased flight speeds in the dark compared to intact-wing controls. Upon exposure to visual contrast in light, the clipped-wing flies reduced their speed comparable to that of the control group flies. Notably, the lower speed persisted upon returning to the dark, indicating an enduring change to the flight controller. To discern between feedforward adaptation and a change in mechanosensory feedback gains, we replicated the experiment after ablating the antennal arista, the primary mechanosensors for sensing airspeed. Although flies with ablated antennae flew with greater variance in speed, they displayed a parallel trend in mean speed adaptation: increased speed in the dark, compensation in the light, and sustained lower speed in subsequent dark conditions. This consistent pattern strongly supports the involvement of adaptive feedforward control rather than the adjustment of mechanosensory feedback gains. Our investigation unveils an adaptive strategy in *D. Melanogaster* flight, illustrating its ability to set flight speed through adaptive feedforward control mechanisms.

## Introduction

Animals must contend with coordinating self-motion despite the ever-changing dynamics of both the neuromuscular system and the external world. Such changes include variations in either the relationship between the external world and physiological sensor responses, or the relationship between physiological motor commands and the resultant forces. In principle, the nervous system could maintain performance under these variable dynamics either by using a robust and unchanging sensorimotor controller or by adapting the controller to the changed dynamics. However, animals throughout the animal kingdom appear to adapt their sensorimotor systems [1, 2]. For example, systematic perturbation of human subjects reaching for a target results in new reaching dynamics that are sustained following further changes in experimental dynamics [3, 4].

In vertebrates, the cerebellum is thought to underly adaptive sensorimotor control through a combination of forward and inverse internal models [5–7]. Forward models compute a prediction of changes in body state and sensation given a desired motor action [6], thereby enabling the nervous system to distinguish between the sensory consequences of self-motion (re-afference) and sensory input arising from external sources (ex-afference) [8]. This distinction prevents stabilisation reflexes from countering volitional movements, whilst allowing the same reflexes to operate continuously to maintain stability. Feedforward commands are estimates of the motor commands required to enact desired movements or, at a minimum, supplement the actuation commands of a feedback controller. This feedforward control avoids many of the sensory delays associated with feedback control [9, 10], as well as being more robust to sensor saturation and noise [10]. Furthermore, by separating re- and ex-afference, the feedforward controller can theoretically be adapted based on an error signal defined by the difference between the expected and actual sensor response, rather than the desired and actual state. This is advantageous as it is not subject to errors that result simply from time delays between control and sensory feedback [11]. Importantly, both forward and inverse models are adaptive filters [9], enabling them to account for the variable dynamics between the external world and the sensory (forward) and motor systems (inverse).

Insects are also adept at adapting their sensorimotor control systems [1, 12, 13]. Insects can compensate for changes in dynamics due to limb loss [14], wing damage resulting from collisions or predation [15–18], mass loading [13, 19, 20], temperature changes [21], and nutritional state [22]. Tethered flying *D. Melanogaster* can orient towards a vertical stripe under closed-loop conditions in which their yaw torque controls the azimuthal position of the stripe, even when the feedback sign between yaw torque and stripe position is inverted [23]. Furthermore, such sensorimotor adaptation is transferable to other motor systems under novel dynamics unlikely to be experienced in nature. For example, flies continue to fixate towards a vertical stripe by using their prothoracic legs to control a lever [1], or using tethered flight yaw torque to turn off a high-temperature harmful laser [23].

Whilst the aforementioned studies indicate that insects can adapt to changes in dynamics, it is not clear whether insects learn to reconfigure a feedback controller, or whether they update a feedforward controller akin to cerebellar inverse models. Evidence for feedforward control has been demonstrated in the fly [24] and wasp [25] gaze stabilization systems; and mantis shrimp strikes [26]; however, whether such feedforward controllers can be adapted in insects remains unknown. Recent behavioural work in *D. Melanogaster* demonstrated that the optomotor feedback controller does not adapt [27]. The frequency and phase responses of the yaw optomotor response of tethered flies in an augmented reality arena were found to be invariant for 30 minutes despite experimental changes in visual feedback gains [27]. This indicates that the insect visual feedback controller is unchanging, thereby implying that adaptive behaviour may indeed arise from adaptive feedforward control.

In this study, we provide further evidence for the existence of feedforward control, and then test its adaptability by controlling the availability of visual feedback following wing damage in *Drosophila melanogaster*. We confirm that changes in flight trajectories due to wing damage can be compensated for in the presence of visual feedback, as previously reported [16]. However, crucially, we find that the compensation for flight speed persists upon returning to the dark, whereas other parameters such as flight curvature and velocity vector elevation angle do not. We show that these lasting changes are still observed after disruption of the antennal mechanosensory system, suggesting that they arise from adaptation of the feedforward component of the flight speed controller, rather than adaptation of the antennal feedback system.

## Methods

### Animals and ablation procedures

A lab-reared colony of recently wild-caught (i.e. within 9 months) *Drosophila melanogaster* fruit flies were maintained on fruit fly media (Advanced Husbandry, Pets and Pieces Ltd., UK) under a 12/12 hr on/off light cycle (8am/8pm). A total of 681 7-28 day old flies were used for the experiments. For clipped-wing conditions (581 flies), flies were cold anaesthetised, and one wing per fly was clipped chordwise from the intersection of the longitudinal veins L1 and L2 (following previous work [28]). For experiments with bilaterally disrupted antennal mechanosensation (241 flies), both antennal aristae were removed from the antennae (following previous work [29]).

### Experimental setup and procedure

Flies were inserted into a 400 *×* 400 *×* 300 mm clear acrylic enclosure with 6 mm wall thickness, via an opaque transfer tube (Figure 1a). An LED strip (white light, 9.6 W m^*−*1^, CRI *>* 80) positioned around the four upper edges of the enclosure provided visible illumination, and a checker pattern was positioned on two side walls to provide visual contrast. The enclosure was continuously illuminated with two infrared lights (96 850 nm wavelength LED, 12 W) diffused with tracing paper directed into the remaining two side walls to permit videography in both light and dark conditions. Two fastcam Sa3 high-speed cameras (Photron Europe Ltd.) with 24 mm lenses (Nikon, AF Nikkor 1:2.8 D, Japan) were positioned beneath the enclosure. The cameras were triggered to capture flight trajectories at 125 fps (aperture F/4, 1 ms exposure time), resulting in a 21.6 seconds (2700 frames) stereo recording every 10 minutes. Each frame was 1024 by 1024 pixels.

**Fig 1.**
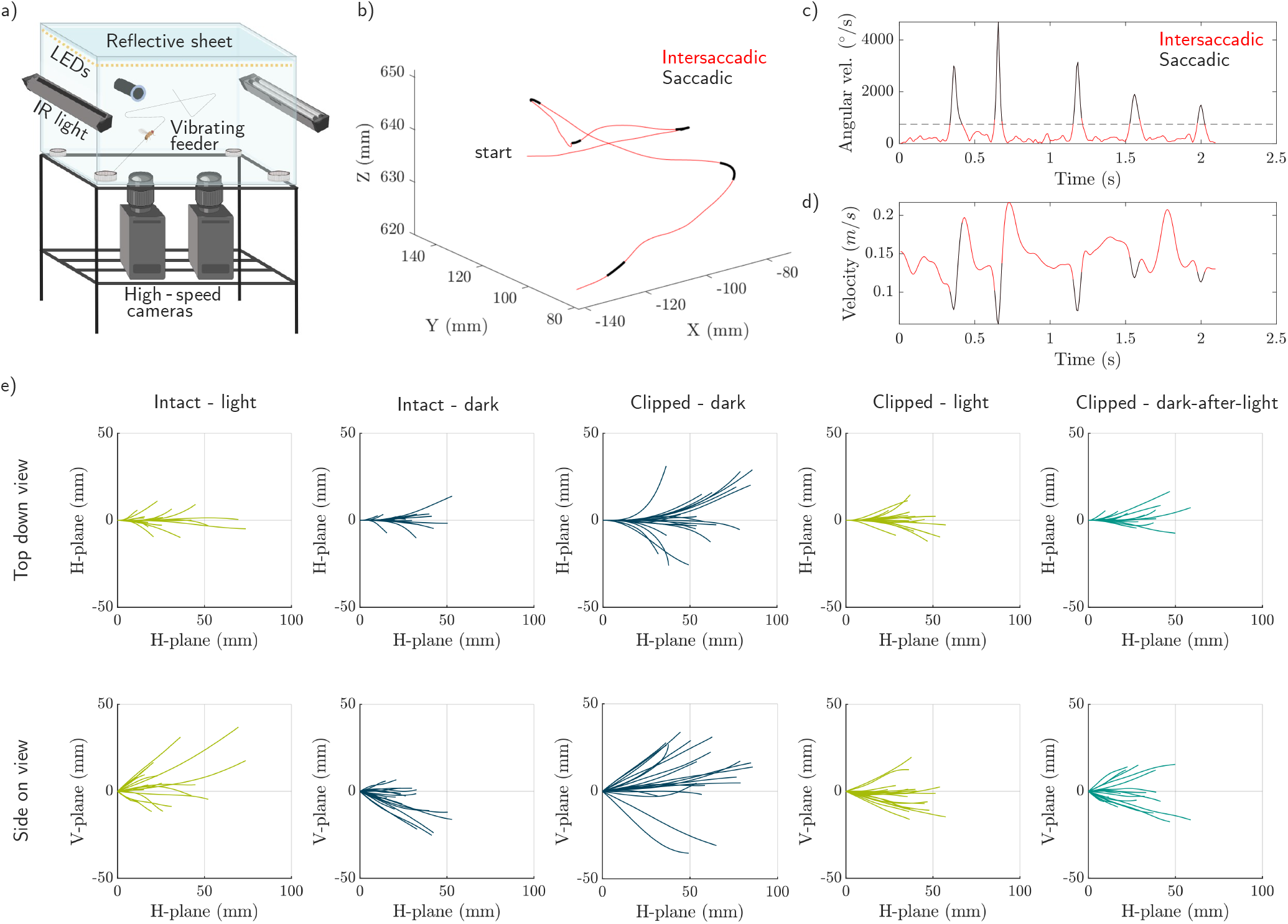
(a) Experimental setup. For clarity we omitted: a checkerboard texture on the near and far enclosure walls; four blackout blankets that kept the entire setup in darkness; wet tissue paper in a cup that provided water and some humidity to the set-up; and the temperature sensor inside the enclosure. The tube on the back wall had a mesh for air circulation and was used for inserting the animals. (b) Trajectory from a fly with intact wings flying in the light (I-L condition). (c) Angular velocity of the velocity vector during trajectory with the threshold angular velocity for a saccade at 750 ^*°*^ s^*−*1^. (d) Translational velocity of the animal. (e) 20 intersaccadic sections of 160 ms from each condition. The intersaccadic sections were rotated to initially start in the same direction and are viewed from the top down, and side on.

Four lures containing single-use disposable tissue soaked in malt vinegar were positioned at each of the lower four corners of the enclosure and were covered by a fine steel mesh so the flies could not access it. Underneath each lure was a Parallax 28822 vibration motor capsule that could vibrate the lure to encourage flight. A custom-designed printed circuit board (PCB) controlled the trigger for the stereo camera setup, the vibrating lures, and the strip of LEDs that went around the top of the enclosure. Before each experiment, a 274-frame calibration (*≈* 12.5 fps) sequence for the stereo-cameras was acquired using a 5 *×* 7 40 mm checkerboard. A temperature sensor (RS PRO-USB-1) was configured to record the temperature every 10 minutes and placed inside the enclosure, along with the vibrating feeders, and damp tissue to provide water and humidity.

We used three visible lighting conditions for wing- and/or aristae-damaged flies throughout the study: LED lights continuously off from the start of the experiment (Dark, D), LED lights on (Light, L), and subsequent dark-after-light (DAL) where LED lights were turned off during video acquisition and on when the trajectories were not being recorded. The DAL condition always followed the L condition.

The eye adapts to luminance levels on the time scale of milliseconds to seconds [30]. To ensure that local luminance adaption or contrast adaption was not a confounding factor in our experiments, during preliminary data collection, we experimented with two dark-after-light (DAL) conditions, (i) lights were off for the entire duration, compared to (ii) lights were only off when recording videos. The only distinction was increased proclivity for flight under the latter condition (see supplementary information S1.2) - hence all data shown here was collected under condition (ii).

For flies with intact wings, we made no distinction between animals in the dark and the dark-after-light since there was no change to the flight dynamics with which they had been reared. Flight trajectories were recorded in the light (4 hrs), then in the dark (4 hrs), then for 4 hrs where LED lights were turned off during video acquisition. This cycle of 12 hrs was repeated to complete one 24 hr period. We initially chose 4 hr intervals to fit within the circadian cycle; however, we did not find noticeable differences in flight propensity or trajectory kinematics for flies subjected to three 8-hour intervals. Therefore, we continued with 8 hr intervals under arista- and/or clipped-wing conditions to maximise the number of trajectories collected per experiment (since the dark/light sequence could not be repeated).

Clipped-wing flies were subjected to dark conditions for 8 hrs, light for 8 hrs, and then DAL for 8 hrs. To exclude nutritional state as a confounding factor a final control experiment was included in which clipped-wing flies underwent a full 24 hrs in the dark.

Finally, the internal body clock of the *D. Melanogaster* has been shown to influence the activity of walking animals [31]. To exclude the possibility of circadian rhythm affecting the results, the start time of the 24 hr experiments was staggered between 9:30 and 18:30. Looking at a subset of data where the three conditions occurred at the same time between midnight and 8 am the trend was consistent, thus excluding circadian rhythm as an explanation for our results (see supplementary information S1.1).

### Tracking and analysis

The first step was the camera calibration of the stereo configuration to allow pixel locations in the two cameras to be transformed into an XYZ position in a global coordinate frame. The points between the checkers were extracted (using the method described in [32]) with a minimum corner metric threshold of 0.2. Using the extracted points, the intrinsic and extrinsic camera parameters were estimated assuming no distortion from the lens, this was followed by a nonlinear least squares minimisation where all parameters were estimated with no assumptions about the lens distortion.

The mean reprojection error of all the calibrations was 0.33 pixels, and the maximum mean reprojection error of any of the calibrations was 0.83 pixels. Following this, a mask was determined that highlighted the tracking area. Then each pair of videos could be run through the tracking algorithm. For each video pair, the relevant mask and camera parameters were loaded. 20 frames equally spaced from the beginning to the end of each video were averaged to generate a background average frame. Each video frame from each camera had the average subtracted and was binarised with a threshold scalar luminance value of 0.05 [33]. Closed areas between 3 and 16 pixels were stored. For each detected object, a Kalman filter was used to determine when consecutive detections were a trajectory by assigning costs based on not being assigned to a trajectory and the distance between the center of consecutive detections. When the trajectories identified on both cameras had significant temporal overlap, they were triangulated for reconstruction in 3D. The 3D trajectories were then saved provided that the root mean square error of the reconstruction was less than 10 mm.

Following the extraction of trajectories, a cubic smoothing spline was applied with a smoothing parameter set to 0.99999 [34]. This parameter was deliberately chosen as it represented the highest degree of smoothing achievable without leading to the underfitting of saccades. A figure demonstrating the tracking process and showing the raw and smoothed trajectory is provided in the supplementary information S2. Trajectories were removed if they were walking, which was determined by speed and proximity to the sides of the enclosure, and removed if they were ascending or descending more than 45^*°*^ from horizontal. A threshold of 750 ^*°*^ s^*−*1^ was applied to the turning rate during each trajectory to determine if the animal was saccading as shown in Figure 1. Finally, intersaccadic flight segments that lasted less than 80 ms were omitted from the analysis. Following these omissions the number of intersaccadic flight segments is given in Table 1.

**Table 1.**
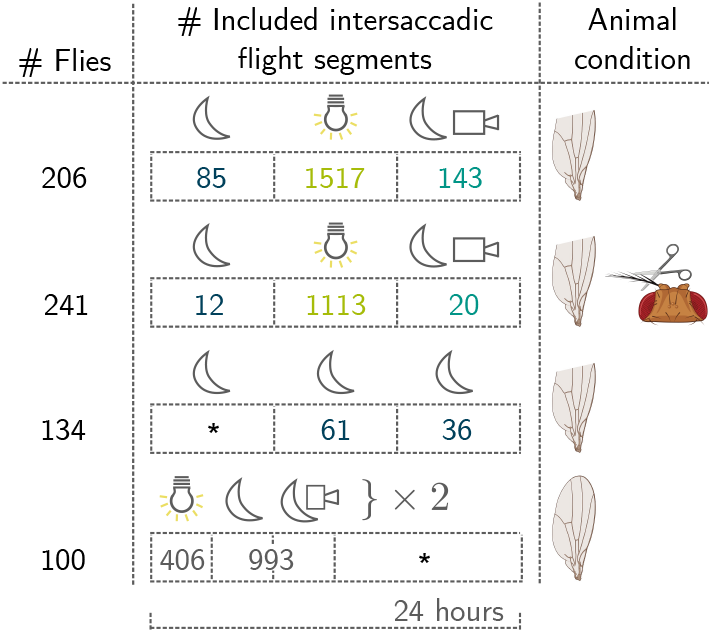
Numbers of animals used in each experimental condition and the number of included intersaccadic flight segments. Sections were only omitted from the analysis if they were: 1) shorter than 80 ms, 2) ascending or descending at an angle greater than *±*45^*°*^, 3) too close to the wall and slow such that they may be walking.

The statistical significances of differences in the intersaccadic kinematics were calculated using a two-sample t-test (Welch’s unequal variance test) where the variance was not assumed to be the same and, therefore, Satterthwaites approximation was used. Tables of results are provided in the supplementary information S3 & S4.

## Results

### Experiment 1: lasting recovery of flight speed following light exposure in clipped-wing flies

Flies with asymmetric wing damage were previously shown to follow curved flight trajectories in the dark, but correct their wing stroke kinematics to maintain straighter flight in the light [28]. We hypothesised that any adaptive changes to the flight controller based on visual feedback would fail to correct flight deviations in the dark due to the lack of input to the motion vision system, but any adaptive changes to a feedforward controller would persist in the dark as this does not rely on visual feedback. We tested this hypothesis by comparing the free-flight kinematics of wing-intact (I-) and clipped-wing (CW-) flies flying in the dark (D), in the light (L), and again in the dark after having flown in the light (dark-after-light, DAL). Intersaccadic flight segments of 160 ms in each condition are shown in Figure 1e.

In agreement with previous work [28], the curvature, a measure of the deviation from a straight line, of inter-saccadic flight trajectories (Figure 2d) was significantly (*P <* 10^*−*4^) larger for clipped-wing flies in the dark (CW-D) (mean *±* std: 4.3 ± 1.9 m^*−*1^), compared with wing-intact flies flying in the light or the dark (3.4 ± 1.3 m, and 3.5 ± 1.3 m^*−*1^, respectively). This increase in curvature in clipped-wing flies was compensated for in the light (2.9 ± 1.3 m^*−*1^), consistent with [28]. However, flies returned to the dark, after flying in the light, in the CW-DAL condition flew with curvature (4.0 ± 1.6 m^*−*1^) insignificantly (*P >* 0.05) different from clipped-wing flies in the dark. This indicates that compensation for inter-saccadic curvature does not persist after returning to the dark, contrary to our hypothesis.

**Fig 2.**
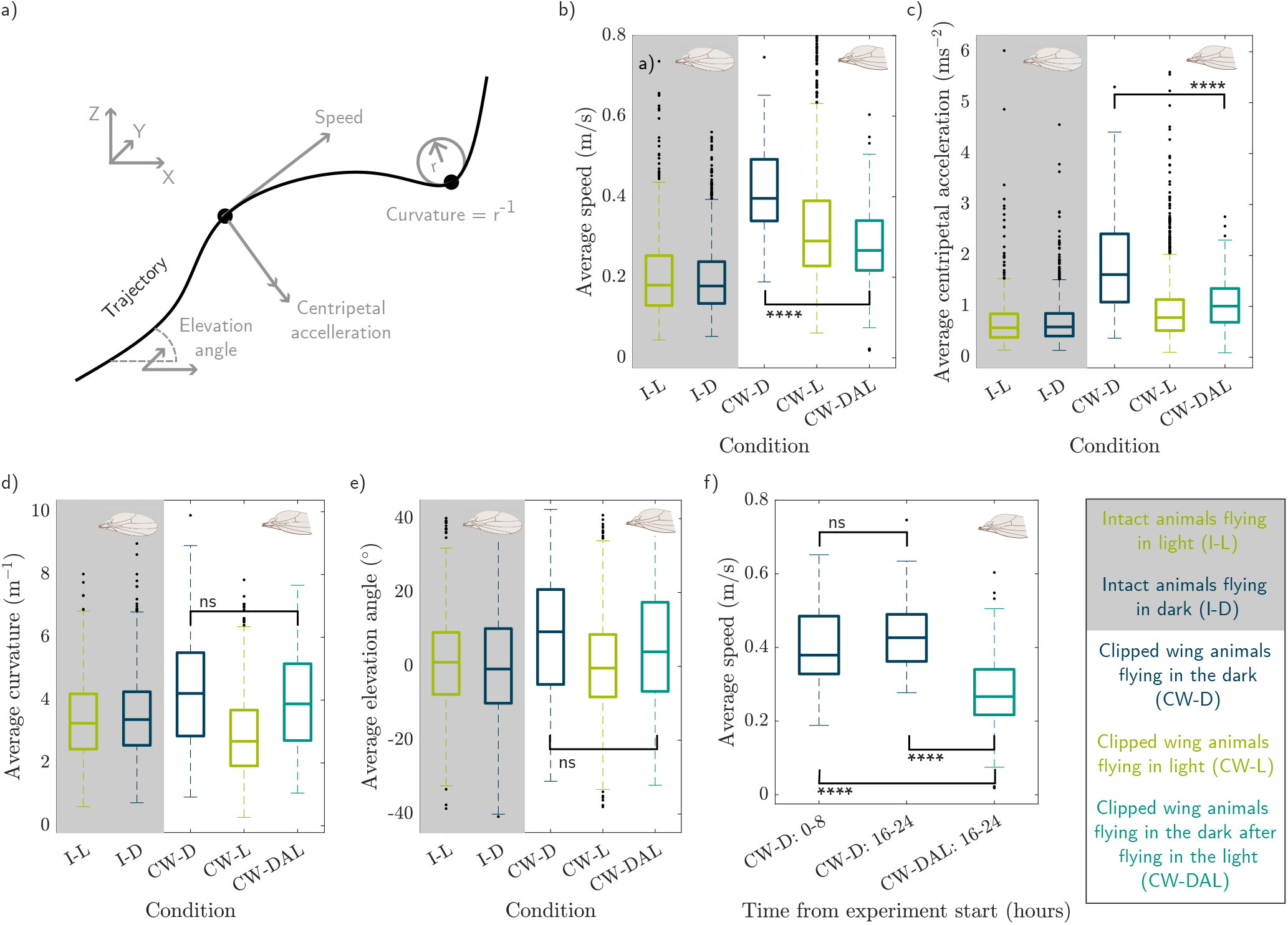
Intersaccadic kinematics for intact and clipped-wing *Drosophila* flying in the light, dark, and dark-after-light. **** indicates *P <* 10^*−*4^, and ns indicates *P >* 0.05. The whiskers indicate the upper and lower quartiles. (a) Is a diagram of a demonstrative trajectory showing the centripetal acceleration, curvature, and speed. All the variables were calculated in 3D. (b) shows average intersaccadic speed for the 5 conditions: intact wings in the light (I-L); intact wings in the dark (I-D); segments from animals with a clipped wing flying in the dark since the wing was clipped (CW-D); clipped wing in the light (CW-L); and clipped wing in the dark, after having flown in the light (CW-DAL). (c), (d), and (e) show the same but for average intersaccadic centripetal acceleration, curvature, and elevation angle respectively. (f) shows the average intersaccadic speed for flies in the CW-D condition for the first 0 - 8 hours of the experiment and the final 16-24 hours of the experiment, next to flies in the CW-DAL condition all from 16-24 hours of the experiment.

However, the curvature describes the resulting geometry of flight trajectories but does not account for the dynamics at which flies progress through these trajectories. Centripetal acceleration is the component of total acceleration normal to the velocity vector (Figure 2a). We found that the centripetal acceleration of clipped-wing flies in the dark was significantly higher (1.8 ± 1.0 m s^*−*2^; *P <* 10^*−*4^) than wing-intact flies flying in either the light or dark (0.7 ± 0.6 and 0.7 ± 0.4 m s^*−*2^, respectively). When the clipped-wing flies were exposed to visual contrast in the light (CW-L) their centripetal acceleration reduced (0.9 ± 0.6 m s^*−*2^) closer to the controls, and significantly lower than in CW-D (*P <* 10^*−*4^). Furthermore, this reduction of centripetal acceleration persisted upon returning to the dark, after flying in the light, (1.1 ± 0.6 m s^*−*2^). CW-DAL condition flies flew with a significantly lower average inter-saccadic centripetal acceleration than CW-D condition flies (*P <* 10^*−*4^). This suggests that whilst flight trajectories geometrically are similarly curved for CW-D and CW-DAL, the centripetal acceleration (and therefore turning force) is lower for the CW-DAL condition than CW-D. Since centripetal acceleration corresponds to velocity squared multiplied by curvature, this indicates that the only substantial persistent change in trajectory kinematics relates to flight speed, rather than curvature.

Indeed, Figure 2b shows the average flight speed of clipped-wing flies in the dark was significantly higher (41 ± 10 cm s^*−*1^; *P <* 10^*−*4^) than wing-intact flies flying in either the light or dark (22 ± 14 and 20 ± 9 cm s^*−*1^, respectively, in line with reported values in [35]). Clipped-wing flies flying in the light (CW-L) were able to compensate for the increased flight speed, flying at a significantly lower flight speed (33 ± 16 cm s^*−*1^) than clipped-wing flies in the dark (*P <* 10^*−*4^) and at a speed closer to wing-intact controls. This reduction of flight speed persisted upon returning to the dark, after flying in the light (CW-DAL) (27 ± 10 cm s^*−*1^). This indicates that compensation for flight speed does persist after returning to the dark, which accounts for the persistent changes observed for centripetal acceleration.

Why do clipped-wing flies fly faster than wing-intact or light-exposed flies? One explanation could be that clipped-wing flies in the dark have ineffective pitch control and lose altitude with gravity, resulting in higher speeds. However, we found that the elevation angle of the velocity vector relative to the xy plane (Figure 2e) was significantly higher for clipped-wing flies flying in the dark (8.3 ± 18 deg; *P <* 10^*−*4^) compared to wing-intact controls (1.3 ± 14 deg, and 0.0 ± 16 deg for light and dark, respectively). Clipped-wing flies therefore flew on average upward against gravity. This elevated angle of the velocity vector was returned to control levels in clipped-wing flies flying in the light. However, this reduction did not persist when they were returned to the dark-after-light; there was no significant difference between clipped-wing flies flying in CW-D or CW-DAL conditions (*P >* 0.05). This suggests that gravity does not explain the persistent change in speed from the dark to the dark-after-light condition.

Another potential explanation for the reduced flight speeds in CW-DAL conditions, compared to CW-D, was that clipped-wing flies reduced their speed as a function of time, either due to fatigue or longer periods of food deprivation. Thus, we compared CW-DAL condition flight speeds with clipped-wing flies flying in the same time window as the CW-DAL condition (16-24 hours) but having experienced only dark for the full 24 hrs (Figure 2f). There was no significant difference in the flight speed of flies in continual dark compared with the initial 8 hrs of dark (*P >* 0.05), whereas flight speed for CW-DAL was statistically significantly different from both the 0-8 and 16-24 hrs dark-only condition (*P <* 10^*−*4^). This suggests that the trend of reduced flight speed in the clipped-wing DAL condition was not a function of time, rather it corresponds to a lasting change in the speed flight controller.

To summarise, there is a clear change in speed from flying in the dark with one wing clipped, to flying in the dark with one clipped after having flown in the light. Clipped-wing flies flying in the dark have a 104 % increase in average speed from wing-intact controls. In the CW-DAL condition, this increase is 35 % from the control group. The adaptation that occurred when visual feedback was available therefore mitigated 67 % the increase in speed caused by the change in system dynamics.

### How could the controller change

How can we account for this persistent reduction in free-flight speed in clipped-wing flies returning to the dark after exposure to the light (CW-DAL)? Figure 3 shows a hypothesised macroscopic model for the insect speed control system [10, 36–39]. The flight dynamics block represents a model for the resulting self-movement following a given motor command. The resulting combination of self-movement and external factors is sensed by the mechanosensory and the visual system. For stabilising behaviours, the mechanosensory and visual control systems sense disturbances, and initiate corrective muscle commands to mitigate course deviation. For goal-directed movements, we hypothesise that a feedforward controller sends the muscle commands to execute the movement; the expected visual sensor response is then cancelled by the forward model.

**Fig 3.**
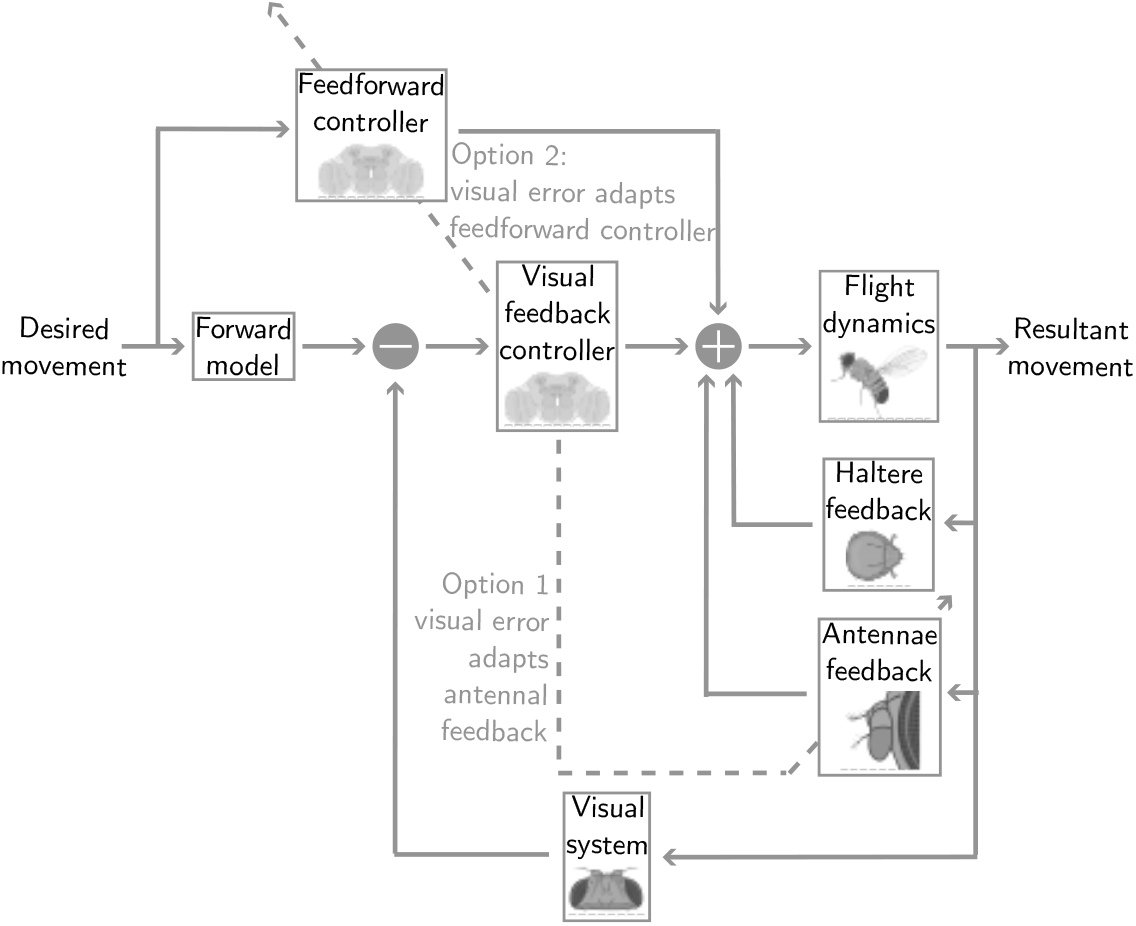
Hypothesised phenomenological model of the insect control system containing visual and mechanosensory stabilising systems in the inner-loop, with a feedforward and forward model component in the outer-loop to accommodate desired movements. Given that we observed a change in speed control, the haltere feedback pathway can be excluded leaving two possible adaptation mechanisms. Option 1 is that the visual error adjusts the antennal feedback system to better control speed when vision is denied. Option 2 is that the induced visual feedback error adapts a feedforward controller - which will also still operate when vision is denied.

#### Option 1: adaptation of mechanosensory feedback

The first of the two plausible adaptation schemes in Figure 3 uses the visual feedback error to adapt the mechanosensory feedback system. Mechanosensation of flight speed is measured by the antennae, which can sense changes in airflow [39]. Tethered flying flies adjust their wingbeat amplitude (and thus thrust) in response to changes in frontal airflow velocity, but this response is absent following antennae removal [40, 41].

However, more recent work examining free-flight behaviour in a wind tunnel where both airspeed and visual scenery could be controlled showed the antennae in *D. Melanogaster* have only a transient effect on flight control [29]. Another mechanosensor thought to be involved in speed control are wind-sensitive hairs. Most of the work on wind-sensitive hairs has been done in *Locusta* [39], and there has been little work in *D. Melanogaster*. However, one study showed that tethered *D. Melanogaster* with their antennae bilaterally deafferented were no better at orienting upwind than dead flies in the same tethered setup [42].

The halteres sense coriolis forces and therefore the angular acceleration/velocity. The halteres are not oriented for measuring translational velocities, and there is no evidence that they are involved in sensing speed [43]. The halteres will be playing a critical role in stabilising flight, and the *D. Melanogaster* would not be able to fly in the dark, with a damaged wing, without the halteres. However, it is unlikely they are responsible for the persistent change in the speed controller observed in Experiment 1.

To summarise, we found no evidence that *D. Melanogaster* have wind-sensitive hairs that are involved in speed control, and halteres do not measure translational velocity. But, antennae have been found to respond to wind speed, where the response was described as transient in [29, 39, 44], and steady-state in [40, 41]. Therefore, it is possible that visual motion information is used to adjust the antennae feedback control system to enhance its performance in case vision is unavailable (option 1, Fig. 3).

#### Option 2: adaptation of feedforward control

The second of the two plausible adaptation schemes in Figure 3 uses the visual feedback error to adapt the feedforward controller. Adaptive feedforward control is well documented in humans [45–47]. In insects, feedforward control has been demonstrated [24, 25, 48], although its capacity for adaptation is hitherto unknown. In the adaptive feedforward model, when clipped-wing animals fly in the dark, they fly faster than intact animals. When exposed to visual contrast in the light, there is an error between the expected and actual optic flow. This is compensated for by the visual feedback controller which acts to reduce the speed. The consistent visual feedback error is used to adapt the feedforward controller until the feedback controller is no longer compensating. Thus, when the flies are returned to the dark, their feedforward muscle commands more accurately reflect the changed system dynamics. As a result, they fly at speeds closer to the control group with intact wings.

### Experiment 2: speed controller changes with bilaterally ablated aristae

Experiment 1 shows that the average intersaccadic speed is significantly different between clipped-wing animals flying in the dark (CW-D), and in the dark-after-light (CW-DAL). After exposure to visual contrast during flight, their speed is much closer to the speed of control flies. There are two possible mechanisms for how the speed control system could have changed to explain the results. The first mechanism is adapting the weights of the mechanosensory feedback system, so that mechanosensors involved in speed control better reflect the changed dynamics from damaging the wing. The second mechanism is adaptive feedforward control.

To distinguish between two potential mechanisms for how the controller could be adapting, the experiments were repeated with animals that had the aristae on their antennae bilaterally ablated, as these are the primary mechanosensor involved in wind-speed sensing.

Figure 4a shows the average intersaccadic speed for animals that have had one clipped wing (CW), and the arista on their antennae bilaterally ablated (ablated antennae, AA). When flying in the dark, the AA-CW-D average speed had a mean *±* std, 89 ± 22 cm s^*−*1^; this is significantly faster than the speed of the control flies (I-D and I-L) presented in the first experiment (*P <* 10^*−*4^). It is also significantly faster than flies in CW-D with wing damage only (41 ± 10 cm s^*−*1^; *P <* 10^*−*4^). The removal of the arista caused an increase in the average speed of clipped-wing animals in D. In addition, there was a significant increase in variance of speed for clipped-wing animals in D when the aristae were removed (Levenes: *P <* 10^*−*4^) consistent with results reported by [29]. When the clipped-wing arista-ablated animals were in the light (AA-CW-L), the visual feedback allowed them to compensate for the increased speed such that they flew significantly slower than AA-CW-D (43 ± 21 cm s^*−*1^; *P <* 10^*−*4^); this was still significantly faster, and significantly more variable than the controls (*P <* 10^*−*4^). When they were returned to the dark, in dark-after-light (AA-CW-DAL), the lower speed persisted (52 ± 32 cm s^*−*1^), and was not significantly different from AA-CW-L (*P >* 0.05), but lower than AA-CW-D (*P <* 10^*−*3^). This implies that the speed controller has changed between AA-CW-D and AA-CW-DAL conditions despite not having aristae.

**Fig 4.**
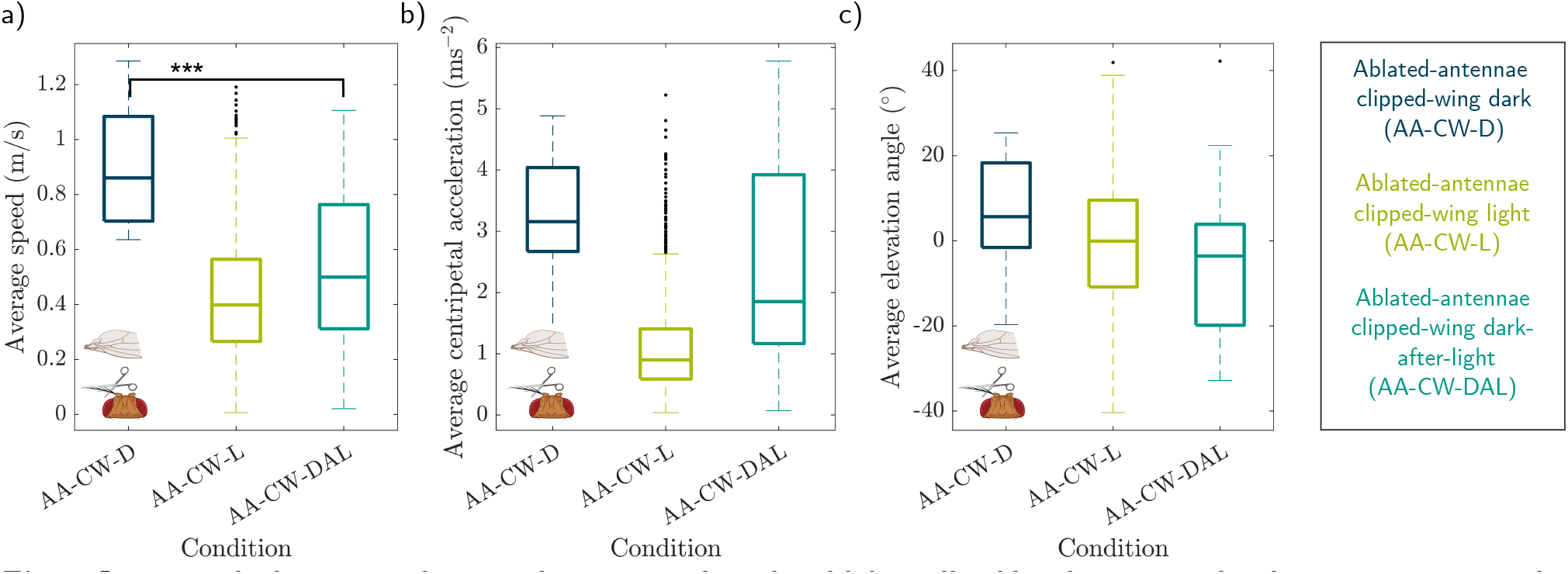
Intersaccadic kinematics for animals one wing clipped and bilaterally ablated aristae under the same experimental paradigm. *** indicates *P <* 10^*−*3^, and the whiskers indicate the upper and lower quartiles. (a) The average inter-saccadic speed for animals flying in the dark (D), then flying in the light (AA-CW-L), and then flying back in the dark (AA-CW-DAL). (b), and (c) are the same for average intersaccadic centripetal acceleration, and elevation angle respectively.

The aristae-ablated and clipped-wing flies had similar trends for centripetal acceleration (Figure 4b) and curvature as those with clipped wings only. In AA-CW-D the centripetal acceleration and curvature were higher than the controls (3.2 ± 1.1 m s^*−*2^ and 3.7 ± 1.4 m^*−*1^, respectively; *P <* 10^*−*4^). These were subsequently reduced when exposed to visual contrast in AA-CW-L (1.1 ± 0.8 m s^*−*2^ and 2.8 ± 1.4 m^*−*1^, respectively). When the flies were returned to dark, AA-CW-DAL, the curvature subsequently increased again to be no different from AA-CW-D (3.7 ± 1.4 m^*−*1^, *P >* 0.05). However, this time the centripetal acceleration for AA-CW-DAL was not significantly lower than AA-CW-D (1.1 ± 0.8 m s^*−*2^, *P >* 0.05).

Again, the increased speed could not be explained by the animals in the dark flying down more. Figure 4c shows that the average elevation angle of the velocity vector for animals in the dark was acting against gravity (7.0 ± 13 deg). This average elevation angle of the velocity vector was slightly higher than in AA-CW-L and in AA-CW-DAL, but not significantly (0.2 ± 15, and −4.5 ± 19 deg, respectively, *P >* 0.05). Thus, the increase in speed in AA-CW-D compared to controls, AA-CW-L, and AA-CW-DAL conditions is not due to falling with gravity.

To summarise, it appears the antennae play a role in both steady-state and transient speed control. When clipped wing animals with and without arista are compared, there are changes in steady-state speed and the variance of speed. Secondly, it appears that without the arista, the animal’s speed control system still adapts. The first experiment shows that adaptation occurs in the speed control system outside of the visual feedback pathway, here, we have shown that it also occurs outside of the antennae feedback pathway. These experiments further suggest that adaptation occurs in a feedforward controller, displayed as option 2 in Figure 3.

## Discussion

In this paper, we examine the ability of *Drosophila melanogaster* to adapt their free-flight kinematics following changes in flight dynamics due to asymmetric wing damage. Experiment 1 found that clipped-wing flies fly faster in the dark compared with wing-intact controls, but that this increase in speed was compensated for when flying in the light. This speed compensation was sustained when light-exposed clipped-wing flies were subsequently returned to the dark (DAL), despite the absence of visual feedback. Overall, the change to the controller under DAL conditions compared to dark mitigated 67% of the speed increase associated with damaging the wing.

We suggested two adaptive control mechanisms that could explain our experimental results (Fig. 3), (i) the animals could use the visual feedback error whilst they are in the light to adjust antennae feedback gains, or (ii) they could use the visual feedback error to adjust a feedforward controller. In a second experiment, we repeated our experimental protocol with flies subjected to both asymmetric wing damage and bilateral removal of antennal aristae, which are the only sensors known to contribute to the control of flight speed in *Drosophila Melanogaster* [29, 39, 40]. Indeed, aristae-ablated clipped-wing flies fly with much higher and more variable speeds than aristae-intact clipped-wing flies, suggesting disregulation of speed mechanosensation, similar to previously reported observations [29]. However, when aristae are removed, the distinct change in speed control from dark to dark-after-light conditions persists to a similar extent as aristae-intact flies. These persistent changes we observed to the flight speed controller in DAL suggests that mechanosensors do not underlie this adaptation.

Interestingly, we found that the compensation of other free-flight trajectory parameters was not sustained upon returning clipped-wing flies to the dark. Both the inter-saccadic trajectory curvature and elevation angle of the velocity vector were significantly higher in clipped-wing flies flying in the dark compared to controls. However, whilst both these kinematic parameters were compensated for in the light, flight trajectories in the dark-after-light were indistinguishable from trajectories in the initial dark condition.

Why do we only find evidence for adaptive feedforward in the control of speed, but not for trajectory curvature or elevation? One of the main advantages of cancelling expected sensor responses with a forward model is to prevent sensor saturation [10]. Neurons have finite output ranges, but by cancelling expected sensor responses to re-afference, any constant response offsets are removed, thus maximising the sensor output range for unanticipated information. When *D. Melanogaster* are flying straight between saccades, any forward model of roll, yaw, and pitch movements should be zero, because the desired trajectory curvature and elevation is zero. However, when controlling for a non-zero speed, the visual sensors will be constantly measuring expansive translational optic flow. To ensure maximum sensitivity and avoid sensor saturation, a forward model would need to cancel this constant translational optic flow. The consequence of such forward model input is that the set-point of any feedback control becomes fixed at zero, i.e. the sensory error available for feedback corresponds only to the difference between expected and actual neuronal response to the optic flow. To actuate a non-zero speed, an additional non-zero feedforward command is therefore required. Adaptation of this feedforward command is what our results suggest accounts for the persistent compensatory reduction in flight speed in DAL.

In the case of trajectory curvature and elevation, since the desired state is zero, a feedforward command is not required. Feedback is therefore sufficient for controlling roll, pitch and yaw. This difference in desired set-point may explain why there were no kinematic differences between dark and DAL in curvature and elevation, whereas these changes were observed for speed. A direct hypothesis that follows from our interpretation is that any flight behaviours following curved and/or elevated trajectories (such as banked turns in *Calliphora*) would follow similar patterns of sustained adaptation as was observed for speed in *Drosophila*.

Our proposed combination of feedforward actuation commands with a forward model cancellation of the expected sensor response for controlling speed (Fig. 3) is consistent with previous results. It is well established that octopamine increases the response gain of optic flow-sensitive interneurons in flies [49–51]. Whilst silencing octopaminergic neurons in *D. Melanogaster* results in no change in average flight speed, corrective accelerations following unexpected optic flow perturbation was lower for octopamine-silenced animals compared to controls [52]. If speed was controlled with a fixed set-point and a feedback controller alone, one would expect octopamine-silenced animals to fly significantly faster than controls, as a larger magnitude of optic flow would be required to sufficiently activate the lower-gain sensors [52]. Alternatively, if average speed is controlled in feedforward (independent of the sensor response) and motion-sensitive interneurons receive forward model input (to cancel the expected sensory consequences of flying with a given speed), then these neurons would only signal unexpected deviations from the expected feedback caused by the feedforward command. This potentially explains why average flight speed is maintained but corrective accelerations are reduced in octopamine-silenced flies [52], because the response of motion-sensitive interneurons to a given unexpected optic flow perturbation is reduced, resulting in a smaller error input to the feedback controller.

The setup was not designed to examine the biomechanics of speed control, however, the results leave two interesting biomechanical questions. Why do they fly faster, and why do they fly up, when in the dark with a clipped wing? The mechanism for controlling speed is done by adjusting the body pitch angle [53]. A higher body pitch angle is associated with a larger wing-damping force, which decelerates the animal [53]. Controlling vertical flight speed is not thought to be done by changing body angle, but instead, wing-stroke [54, 55]. Thus, these two observations are not contradictory and are likely a combination of the wing-stroke and body angle.

Lastly, adaptive feedforward control, rather than adaptive feedback, accounts for the discrepancy that goal-directed behaviours in insects can be highly adaptive [12], but stabilising reflexes are not [27]. This is exemplified by the observation that insects can cope with an inversion of dynamics for orienting towards a bar [1], but were observed to spin uncontrollably for over an hour when wide-field background motion was inverted [56]. When a forward model cancels expected sensor responses from goal-directed behaviours a feedforward controller must correctly actuate the corresponding movements. In contrast, stabilisation reflexes by definition can only be executed with feedback control. The combination of adaptive feedforward control and inflexible feedback control therefore explains long-standing observations that goal-directed behaviours but not stabilisation reflexes are adaptive.

## Supporting information

### S1.1 Circadian rhythm

The internal body clock of the *D. Melanogaster* has been shown to influence the activity in walking animals. To exclude this as a confounding factor in our experiments, the start times of the 24 hr experiments were staggered between 9:30 and 18:30. Figure S1a shows the intersaccadic flight speeds on clipped wing animals in each of the lighting conditions at different times of day. At all times of the day when there were CW-DAL observations, the speed was lower than that of CW-D at the same time of day, suggesting that the results are not due to circadian cycles. Figure S1b further supports this; there was data for each of the three lighting conditions between the hours of midnight and 8 am, and the trend was consistent with that in the manuscript: higher speed in CW-D and lower speed in CW-L and CW-DAL.

### S1.2 Light adaptation state

As stated in the manuscript: during preliminary data collection, we observed consistent trends in the dark-after-light (DAL) condition, where (i) lights were off for the entire experiment duration, compared to (ii) lights were only off when recording videos. The only distinction was increased proclivity for flight under the latter condition - hence all data shown in the manuscript was collected under condition (ii).

Figure S1c shows the preliminary data in which the DAL data was collected under condition (i), plotted next to D and DAL in the manuscript collected under condition (ii). There was no difference in DAL between the two conditions (*P >* 0.05), but a significant difference between D and DAL for both recording conditions (*P <* 10^*−*4^, for both). This excludes the possibility of contrast gain or light adaption state as a possible explanation. Additionally, Figure S1d shows condition (i) and condition (ii) for the 4 hr intervals used for intact animals flying in the dark, again there is no significant difference in flight speed between the two suggesting the light adaption state has no impact on flight speed.

**Table S3.**
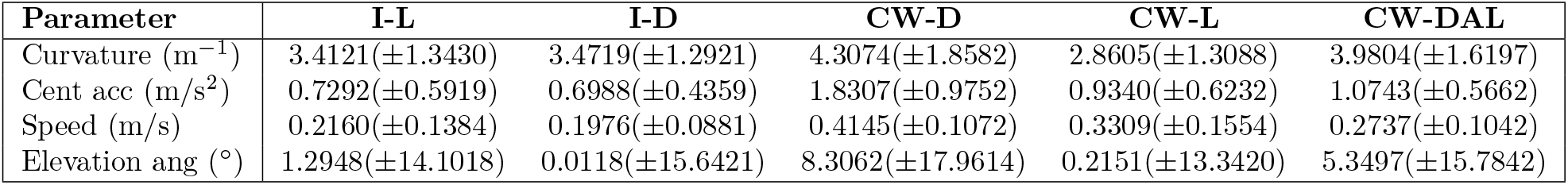
Table of mean and standard deviation of data in experiment 1.

| Parameter | I-L | I-D | CW-D | CW-L | CW-DAL |
| --- | --- | --- | --- | --- | --- |
| Curvature ($\text{m}^{-1}$) | 3.4121($\pm 1.3430$) | 3.4719($\pm 1.2921$) | 4.3074($\pm 1.8582$) | 2.8605($\pm 1.3088$) | 3.9804($\pm 1.6197$) |
| Cent acc ($\text{m/s}^2$) | 0.7292($\pm 0.5919$) | 0.6988($\pm 0.4359$) | 1.8307($\pm 0.9752$) | 0.9340($\pm 0.6232$) | 1.0743($\pm 0.5662$) |
| Speed (m/s) | 0.2160($\pm 0.1384$) | 0.1976($\pm 0.0881$) | 0.4145($\pm 0.1072$) | 0.3309($\pm 0.1554$) | 0.2737($\pm 0.1042$) |
| Elevation ang ($^\circ$) | 1.2948($\pm 14.1018$) | 0.0118($\pm 15.6421$) | 8.3062($\pm 17.9614$) | 0.2151($\pm 13.3420$) | 5.3497($\pm 15.7842$) |

**Table S4.** Table of mean and standard deviation of data in experiment 2.

| Parameter | AA-CW-D | AA-CW-L | AA-CW-DAL |
| --- | --- | --- | --- |
| Curvature ($\text{m}^{-1}$) | 3.7425($\pm 1.3811$) | 2.8122($\pm 1.3813$) | 4.5371($\pm 1.9442$) |
| Cent acc ($\text{m/s}^2$) | 3.2396($\pm 1.0699$) | 1.1208($\pm 0.7727$) | 2.3569($\pm 1.7969$) |
| Speed (m/s) | 0.8902($\pm 0.2221$) | 0.4255($\pm 0.2146$) | 0.5199($\pm 0.3207$) |
| Elevation ang ($^\circ$) | 7.0102($\pm 13.0224$) | 0.2489($\pm 14.9748$) | -4.4830($\pm 19.0975$) |

**Fig. S1.**
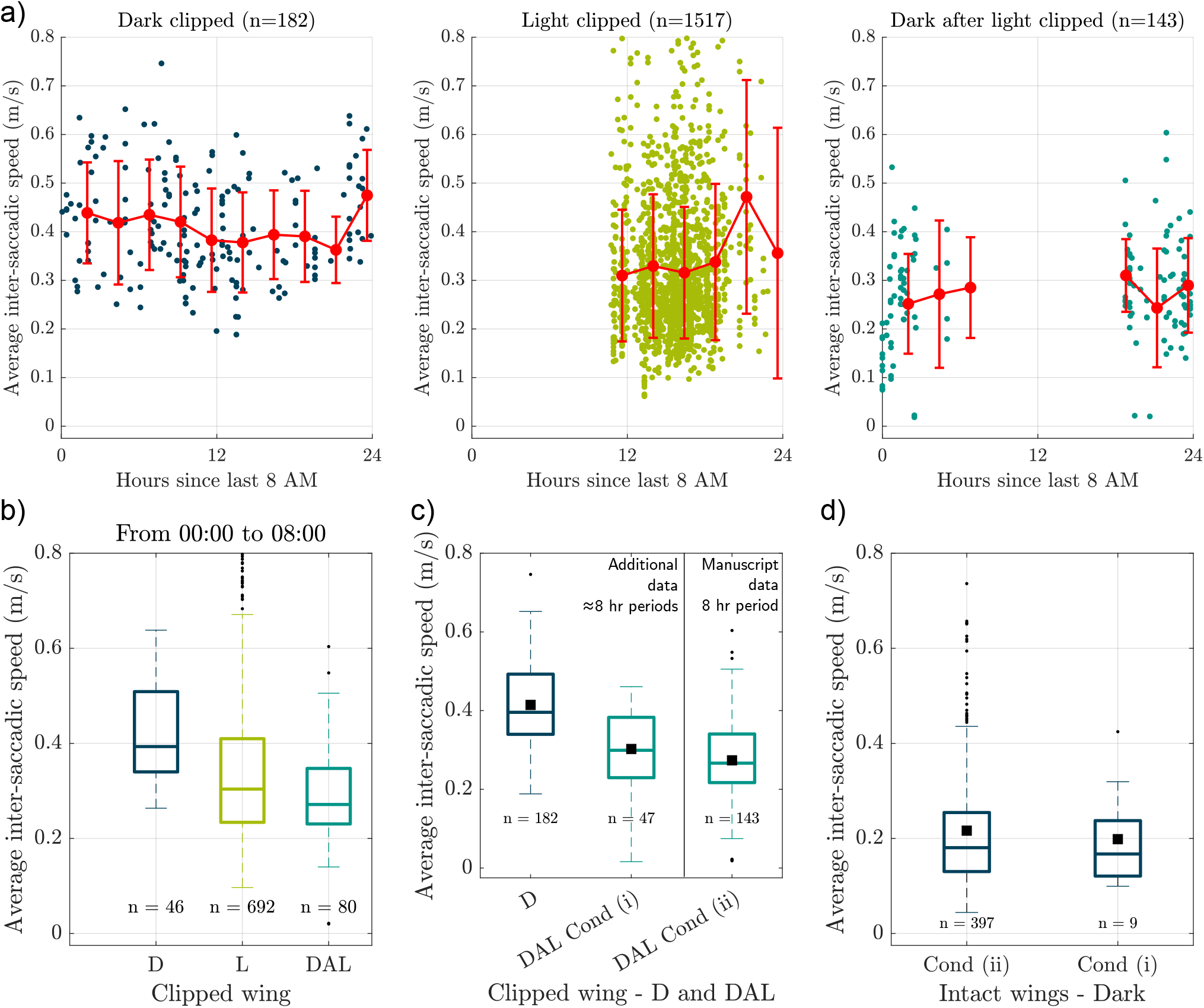
(a) average intersaccadic speed at different times for clipped wing animals in each lighting condition in the manuscript. (b) Shows the average inter-saccadic speeds for observations that all occurred between midnight and 8 am. (c) The average intersaccadic velocity of wing-damaged animals flying in the dark (CW-D); and in the dark-after-light (CW-DAL). Condition (i) is the lights off the whole time, and condition (ii) is they are only off when recording videos. (d) Condition (i) and (ii) for the intact animals in the dark.

**Fig. S2.**
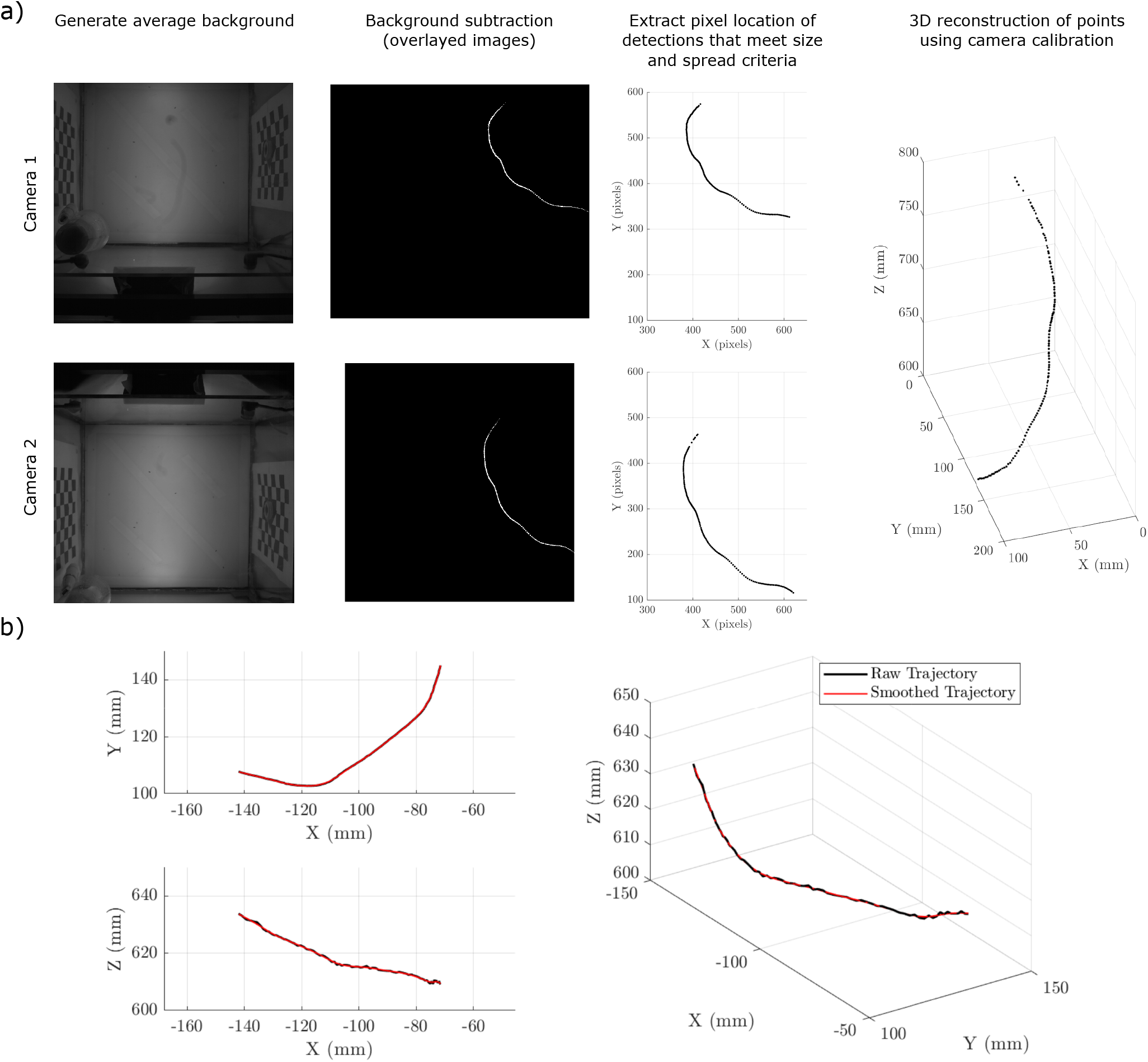
(a) The tracking and reconstruction procedure with an example trajectory. (b) Smoothed and raw trajectories, due to the camera placement the XY axis reconstruction contained very little noise. The Z-axis contained more noise but the difference between reconstructed and smooth rarely exceeded 2.5 mm.

## Acknowledgments

We would like to thank Glenn D’Mello for the helpful and enjoyable discussions on the work. Aspects of Figures: 1, 2, 3, and 4; and Table 1 were made with biorender.com. This work was supported by the Defence Science and Technology Laboratory (DSTLX1000161145).

